# Prefrontal neuronal dynamics in the absence of task execution

**DOI:** 10.1101/2022.09.16.508324

**Authors:** Shusen Pu, Wenhao Dang, Xue-Lian Qi, Christos Constantinidis

## Abstract

Prefrontal cortical activity represents stimuli in working memory tasks in a low-dimensional manifold that transforms over the course of a trial. Such transformations reflect specific cognitive operations, so that, for example, the rotation of stimulus representations is thought to reduce interference by distractor stimuli. Here we show that rotations occur in the low-dimensional activity space of prefrontal neurons in naïve monkeys, while passively viewing familiar stimuli. Moreover, some aspects of these rotations remain remarkably unchanged after training to perform working memory tasks. Significant training effects are still present in population dynamics, which further distinguish correct and error trials during task execution. Our results reveal automatic functions of prefrontal neural circuits, allow transformations that may aid cognitive flexibility.

## INTRODUCTION

Neurons in the prefrontal cortex (PFC) represent sensory stimuli in a dynamic, task-specific manner ^1, 2, 3^. Analysis of population activity with methods of dimensionality reduction reveals that stimuli are typically represented in a low dimensional space, or manifold, with the majority of the firing rate variance captured by a few dimensions ^4, 5^. Furthermore, the representation of stimuli in the reduced space changes dynamically during the course of a trial, as the subjects perform a cognitive operation to adapt to task demands; the stimulus manifold may therefore be projected to a different subspace, rotated, or otherwise geometrically transformed ^6, 7, 8^. Orthogonal rotation of a stimulus representation has been proposed as a possible mechanism for reducing interference between sensory and memory representations, protecting the memory of an initial stimulus from the interference of a subsequent stimulus presentation ^9^. The same stimuli are represented in different subspaces when used in the context of different tasks ^10^ and errors are characterized by changes in stimulus representation geometry ^11^.

While it is clear that geometric transformation of stimulus information in neuronal populations occurs during cognitive operations ^12^, less is known on how the acquisition of a cognitive task may alter stimulus representation geometry. We thus addressed this question by analyzing prefrontal populations, both before and after subjects were trained to perform working memory tasks involving identical stimuli. We examined multiple aspects of the activity space, including subspace alignment, geometrical similarity, and dynamics, to determine the relative contribution of training and regional specificity in the formation of the observed low dimensional geometry.

## RESULTS

Neurophysiological recordings were collected from the lateral PFC of six monkeys in total; pre-training data were acquired from all six animals, and post-training data were recorded from three of these six subjects (Table S1)^13, 14^.Once fully trained, the monkeys viewed two stimuli appearing in sequence with intervening delay periods between them and reported whether the second stimulus (sample) was the same as the first stimulus (cue) and constituted a match, or was different and constituted a nonmatch (Fig. 1A-B). The stimulus sets used in these experiments varied in terms of spatial location or shape (Fig. 1A-B). Recordings were also obtained from these animals, viewing the same stimuli presented with the same timing prior to any training in the task. A total of 1164 neurons from six monkeys in five prefrontal subdivisions (Fig. 1C) were recorded during the passive, pre-training viewing of spatial stimuli; 847 neurons were recorded during the passive, pre-training viewing of feature stimuli. Additionally, 1031 neurons from three monkeys were obtained after training, when the animals performed actively the spatial working memory task; 796 neurons were obtained during the active feature working memory task (see Supplementary Table S1).

**Figure 1.**
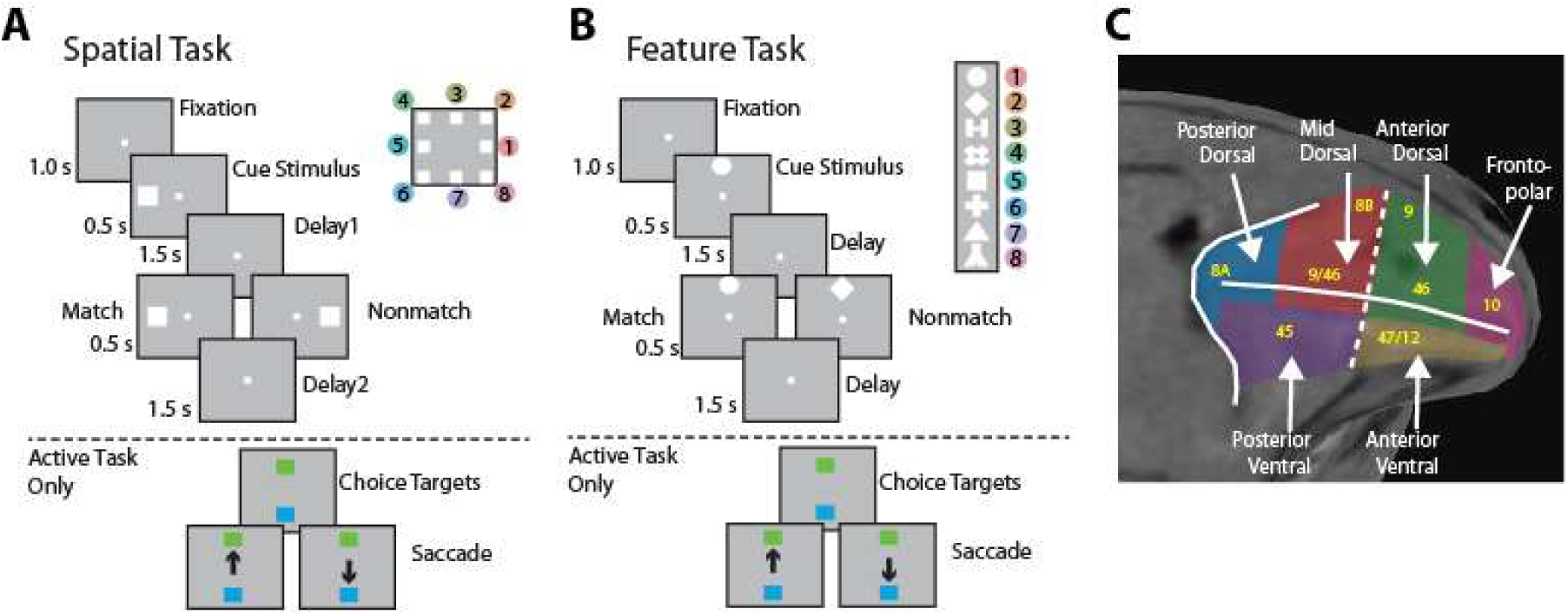
Task structure and recording areas. The animals were required to maintain center fixation throughout both active and passive task trials. At the end of active tasks trials, monkeys were required to make a saccade to a green target if the stimuli matched or to a blue target if the stimuli did not match. (A) Spatial match-to-sample task; the eight cue locations analyzed are shown in the inset. (B) Shape feature match-to-sample task; eight possible shapes in a session shown are shown in the inset. (C). Neurons were recorded from 5 subdivisions of lateral PFC, which we refer to as Anterior Dorsal (AD), Anterior Ventral (AV), Mid-Dorsal (MD), Posterior Dorsal (PD), and Posterior Ventral (PV).

### Spatial stimuli are geometrically arranged in a low-dimensional subspace

We applied Principal Component Analysis (PCA) on the covariance matrix of the zero-centered data, and selected eigenvectors sorted in decreasing order in terms of explained variance. In agreement with previous studies suggesting that prefrontal neurons represent stimuli in a low-dimensional manifold ^4, 5^, we found that, on average, 73% of the variance across brain regions and training phases could be explained by the first three principal components across brain regions and training phases (Fig. S1A and Table S2). This result validated our approach of visualizing and measuring geometry structure in low dimensional space. Our analysis of plane rotations was replicated in higher-order subspaces as well (Fig. S1B-C), with qualitatively similar results.

Previous studies have also revealed that spatial information is arranged in a well-structured geometry resembling the physical appearance of visual stimuli in working memory ^6, 15^, however, it is not clear whether task training may be responsible for the development of this geometry. Our analysis revealed that spatial stimuli (Fig. 2) exhibited a ring-like pattern. This observation was made within the subspace defined by the first two principal components, which preserved the relative position of stimuli in the low-dimensional subspace (Fig. 2A). In contrast, shape stimuli did not demonstrate this pattern (see Fig. S10).

**Figure 2.**
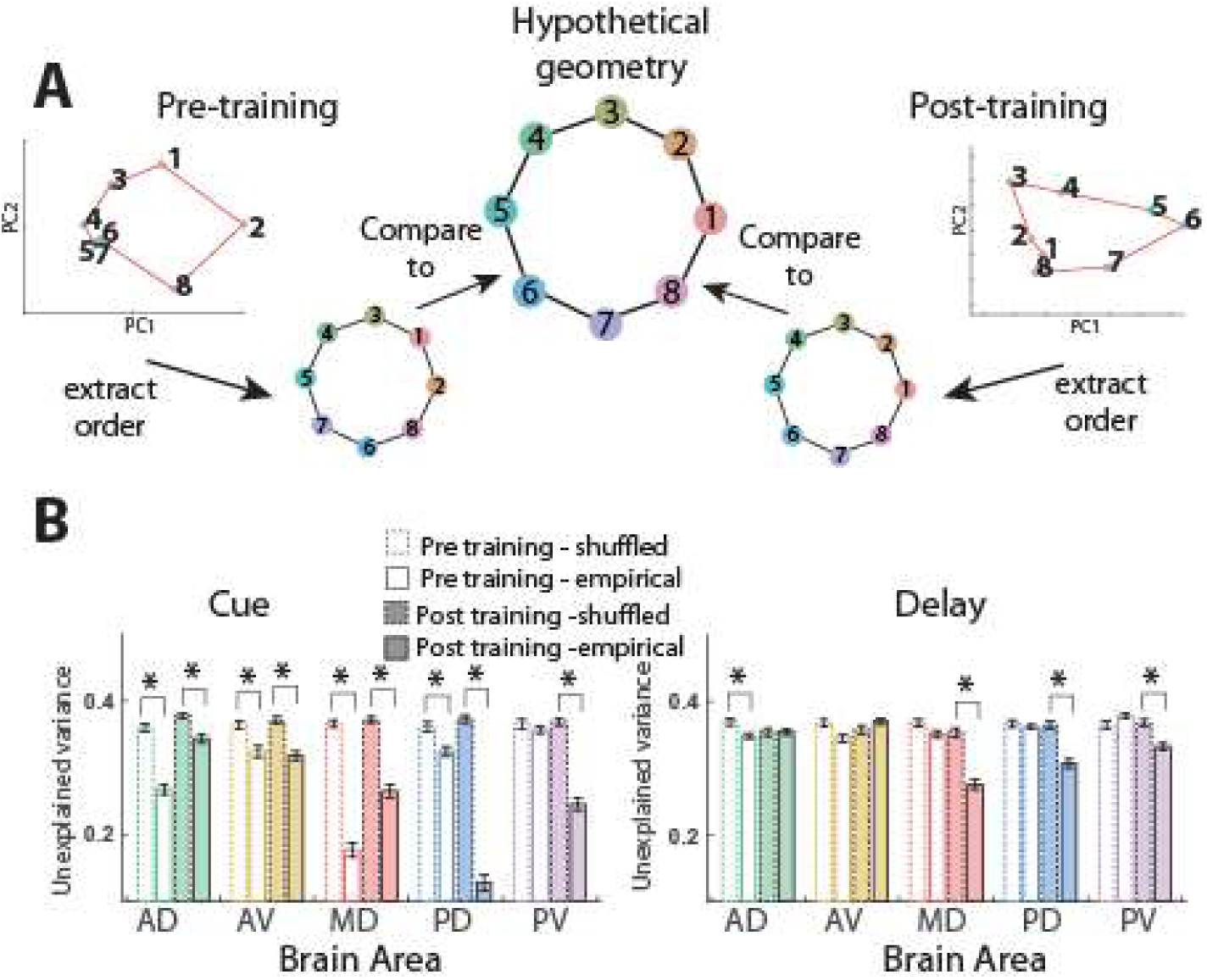
Comparison of within-subspace geometry for the spatial task. (A) Graphical summary of how representational geometry order was compared. Geometries were calculated from cue period activity of the spatial task, pre- and post-training. The geometries were compared to a hypothetical geometry that maximizes the distance between match and nonmatch stimulus pairs (see Methods: Geometry of low-dimensional representation). (B) Training increased the orderly structure in the delay period of the spatial task. Geometrical similarity to the hypothetical configuration for both tasks in the cue and the first delay period was quantified by the unexplained variance after fitting to a standard geometry. To quantify the contribution of the factor of representation arrangement order (e.g. 1, 2, 3, 4 vs. 3, 2, 1, 4 on a ring) independent of the factor of geometrical shape (deviation from vertices of an octagon), the difference between the empirical and an identity-label-shuffled dataset was compared. Color filled bars represent post-training data, empty bars represent pre-training data. Solid line bars represent empirical data and dashed line bars represent the similarity measure after shuffling shape labels, as a control. A larger difference between the filled and the dashed bars indicates a higher order similarity between the standard hypothetical geometry order and the empirical measurement. Test statistics for difference between this order similarity from the pre- and post-training phase in Table S3. Statistically significant differences are indicated by an asterisk (*) above the corresponding bars.

While no task training was required for the establishment of this geometric structure, we investigated if training could enhance its order regularity. More specifically, we explored whether training could increase the order similarity between the empirical geometry order of the eight stimulus conditions and the natural order of eight vertices in an octagon (Fig. 2A). To quantify this similarity, we calculated the unexplained variance between the empirical geometry order and the ‘standard’ order that maximizes the distance between diametric locations. The order regularity was defined by the difference in the unexplained variance between the empirical dataset and an order shuffled dataset. By comparing different prefrontal subdivisions, we found that for the cue period, the anterior and middle regions of PFC already exhibited a strong alignment with the ‘standard’ order before training, while for the first delay period all areas showed little to no order regularity before the animals were trained for working memory tasks. We also found that in general, training increased the “order regularity” for mid-posterior regions, for both the cue and the delay period. This was most pronounced for the delay period in the mid-dorsal and posterior-dorsal subdivisions (Fig. 2B, nonparametric bootstrap test, for pre-vs. post-training order regularity, MD, PD, and PV, p<0.001 in each case).

### Some subspace transformations are task independent

Since low-dimensional stimulus representations in neuronal population activity are transformed during cognitive operations ^7, 8, 9^, we sought to test how such transformations differ systematically before and after training to perform a cognitive task. Previous research suggested that the rotation under different contexts is beneficial for the task, since when two subspaces were orthogonal variation in one subspace will have near zero variation in the other (Fig. 3A2). For example, when one plane in a 3-dimentional space is viewed from the space spanned by another plane, it would have a much larger projection if the two planes are parallel to each other, whereas if they are orthogonal the first plane would only project as a line to the other. We thus quantified transformations based on the rotation of subspaces in different epochs. This was done by measuring the primary angle between the low-dimensional subspaces that accounted for the most variance in different contexts (Fig. 3A1). Measured rotation angles were compared to a baseline rotation, to control for the difference in response variation and selectivity for various PFC subdivisions. This baseline rotation was measured by calculating the rotation between the empirical dataset, and a synthetic dataset (see Methods: Dimensionality reduction and rotation of subspaces) that aim to mimic the statistics of the empirical one.

**Figure 3.**
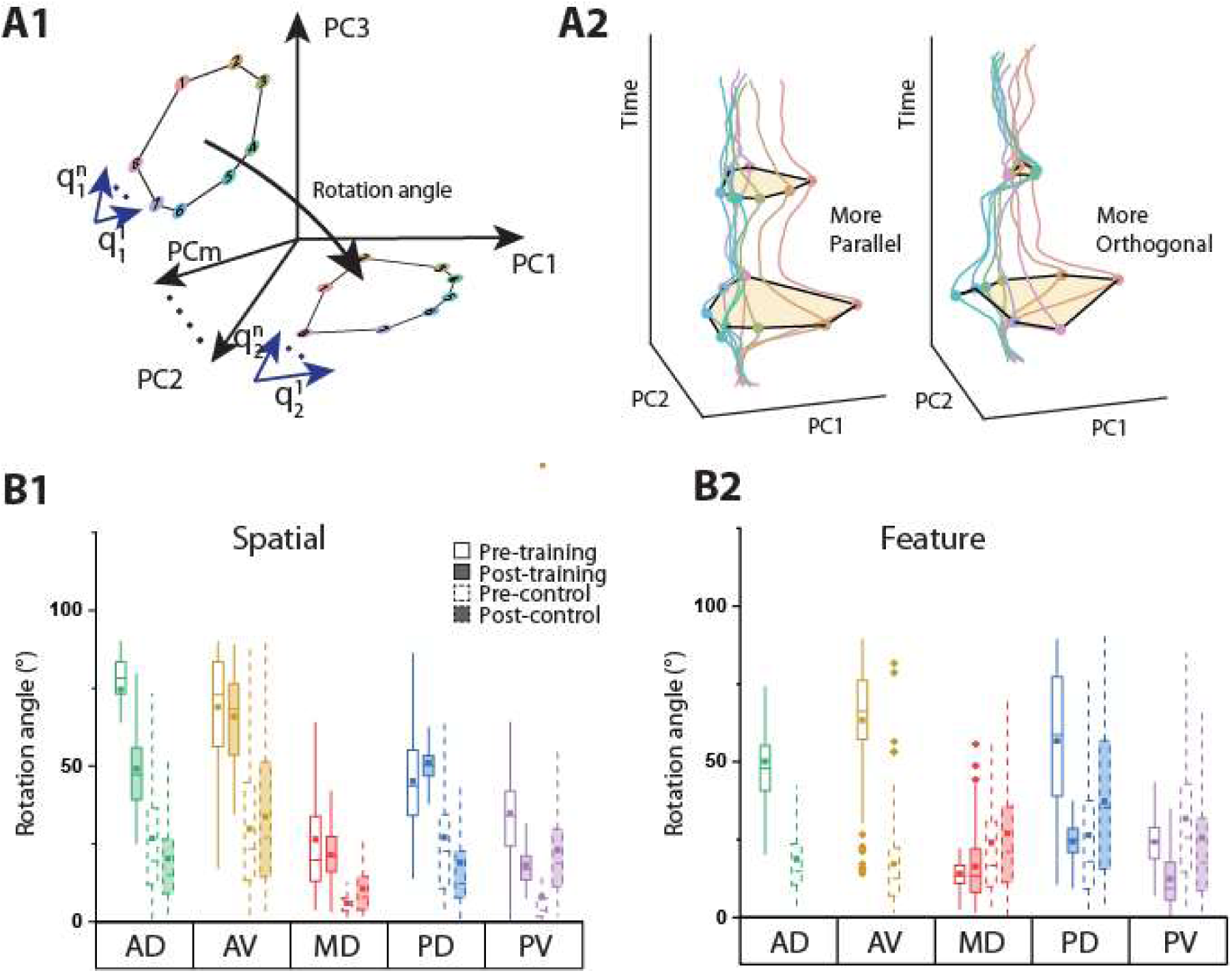
Subspace rotation between the cue and match epochs. (A1) Illustration of subspace rotations. Neural response of different conditions were projected onto a m-dimensional PCA-space (m=3 in Fig. 3-5, m=3-15 in Fig. S1), where the rotation and geometrical similarity were calculated (see Methods: Geometry of low-dimensional representation for details). (A2) Neural trajectory projected in the cue subspace when two representations were more parallel (MD cue vs. match) vs. more orthogonal (MD cue vs. nonmatch). When the second representation was parallel (left) to the first one, the projection on the first subspace was larger, compared to the condition of two representations being orthogonal (right). (B) Solid line bars: angles between the cue and the match condition in different PFC subdivisions pre-(empty boxes) and post-training (color filled boxes) for the spatial (B1) and the feature task (B2). Dashed line bars (control): angles from the control data obtained by generating surrogate data. Dot and horizontal line in the box plot represent the mean and median, respectively; The bottom and top edge of each box are the 25th and 75th percentiles of the sample, and the whisker represents 1.5 times the interquartile range. Pre vs post nonparametric test: spatial AD p=0.369; spatial AV p=0.663; spatial MD p=0.946; spatial PD p=0.909; spatial PV p=0.535; feature MD p=0.440; feature PD p=0.164; feature PV p=0.216. All empirical angles were significantly (p<0.001) larger than the corresponding control angle measurements (Table S4), except for the post-training stages in the PV subdivision.

We examined specifically the geometry of our stimulus set in neuronal activity during the cue presentation and the delay period that followed it; during the cue and match presentations; and during the match and nonmatch presentations. (see “Dimensionality reduction and rotation of subspaces” in Methods).

Strikingly, even before any task training, representations of the same stimuli during the cue and match period already exhibited significant rotation angles in multiple brain areas, for both the spatial and feature sets (Fig. 3B1-B2). Our analysis further revealed that the pattern of transformation between the cue and match stimuli was highly area-specific. In other words, rotation angles differed across regions, while similar angles were seen in the same subdivision before and after training (Fig. 3B1-B2). The Pearson correlation coefficient between pre- and post-training rotation angles was r=0.83 (p=0.079) for the spatial dataset (Fig. S2 left). Similar to the cue vs match trials, we found that substantial rotations between the cue and nonmatch representations already were evident in naïve animals (Fig. S3B), and the rotation angles remained quite consistent across different subdivisions and training status.

To further substantiate the subspace orthogonality observed in the angle measure, we calculated the variance accounted for (VAF) ratio between the cue and the match period (Fig. S4A-B, top panel). The VAF, with a value range from 0 to 1, is a measurement of subspace alignment. Higher VAF values indicate better alignment, while VAF values close to 0 indicate orthogonality (see Methods: variance accounted for ratio - VAF). The results in Fig. S4 qualitatively agree with the analysis of rotation angles (Fig. 3 and 4), with higher angle measurements consistently corresponding to lower VAF ratios. Once again, we found that the relative order of values across subdivisions was preserved between the pre- and post-training phases. This result indicates that populations of neurons in different subregions of the prefrontal cortex transform matching stimuli in a stereotypical fashion, independent of training and task execution.

**Figure 4.**
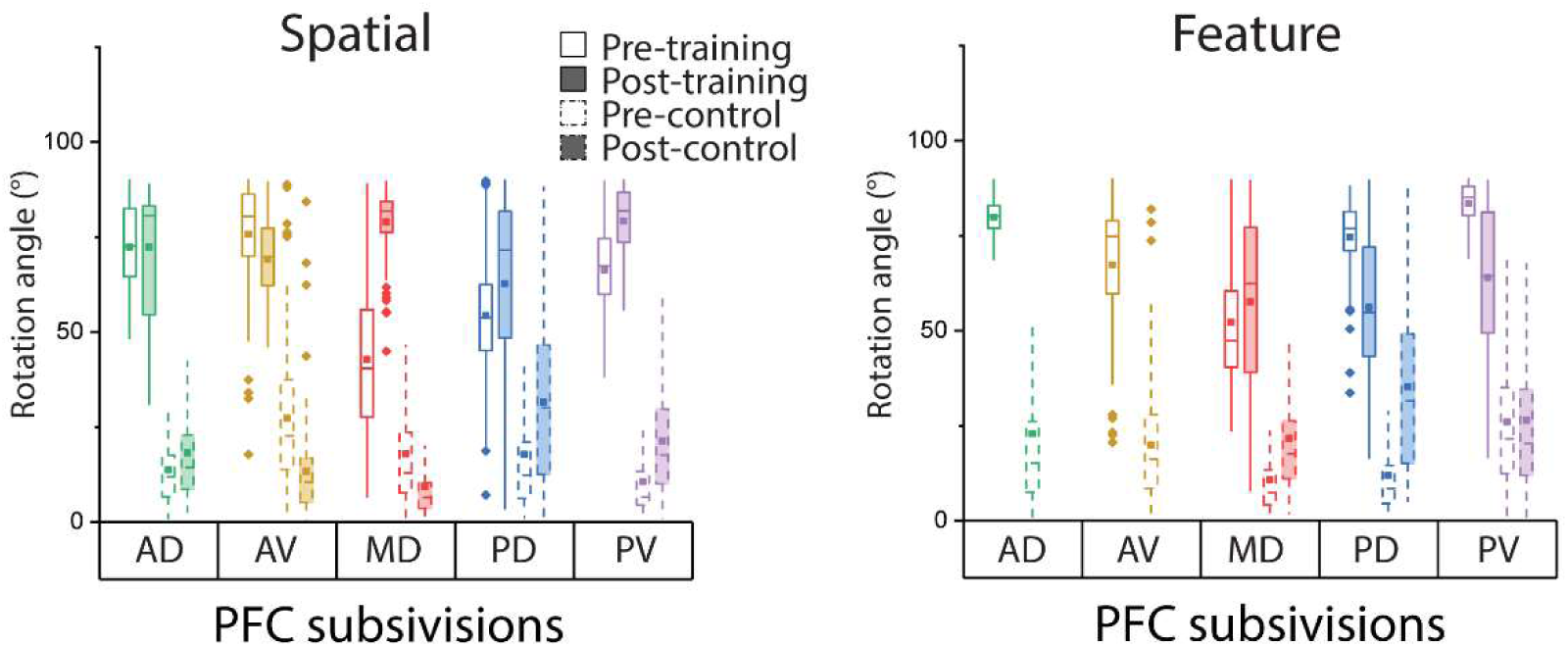
Subspace rotation between the match and nonmatch conditions. Primary angles between the match and nonmatch in different PFC subdivisions for the spatial and the feature task. (pre vs post nonparametric test: spatial AD p=0.385; spatial AV p=0.380; spatial MD p=0.002; spatial PD p=0.569; spatial PV p=0.096; feature MD p=0.090; feature PD p=0.343; feature PV p=0.546). The bottom and top edge of each box are the 25th and 75th percentiles of the sample, and the whisker represents 1.5 times the interquartile range. As a control we randomly drew data from a particular task epoch (empirical: solid bars; control: dashed bars). In this comparison, the control angles were measured by fitting a Poisson distribution to the firing rates in the match trials. All empirical angles were significantly (p<0.001) larger than the corresponding control angle measurements (Table S5).

We were also interested in how single-cell properties contributed to the rotation phenomenon observed at the population level. We examined repetition suppression, the phenomenon of decreased response to a stimulus that is repeated in a trial (match, in the context of our task) over a stimulus that is not repeated (nonmatch) ^16^. This turned out not to have a substantial impact on population rotation measurements; removing the most suppressed cells still yielded results that were highly correlated with the original dataset (Fig. S5). Similarly, we investigated whether the number of cells contributing to the low-dimensional subspace that accounted for most variance (participation ratio) changed across different contexts, or if the tuning of individual neurons to stimuli changed across different contexts. Our analysis showed that the former, difference in participation ratio in various contexts, better explained the rotation angle: areas with highest rotation angles typically exhibited the greatest difference in participation ratio (Fig. S6B-C top).

Although we have emphasized so far the task-independent nature of rotations, some representation transformations were only observed after training, for example involving the representation of stimuli in match and nonmatch conditions. In the spatial task, the angle of rotation between the match and nonmatch representations changed considerably after training (Fig. 4, left) in the most spatially selective, mid-dorsal region^17^. Consequently, across areas, there was little correlation between the plane angles observed before and after training across areas, with the correlation coefficient being r=-0.29 (p=0.491, Fig. S2 middle). The VAF ratio measurement agreed with angle measurement in this case too, evidenced by a major decrease for the mid-dorsal region in the spatial task (Fig. S4-A middle). Similar training effects were observed for the subspace angle between the cue and the first delay period, with anterior prefrontal areas exhibiting increased rotation, particularly for the spatial task (Fig. S3A).

We also compared the dynamic trajectory of population activity before and after training. As expected, pre-training activity was largely confined to a narrow subspace during the delay period, while post-training activity became increasingly dynamic in an expanded space (Fig. S7A, B). To further quantify this change, we randomly sampled half of the trials from all periods and calculated PCA space as the base to project the remaining trials (see Methods: Dimensionality reduction and rotation of subspaces). The mean ratios of the decoding space area between fixation, first delay and second delay period (using the fixation period as reference) were 1 : 1.07 : 1.26 for the pre-training data and 1 : 8.62 : 1.83 for the post-training data (Fig. S7C-E).

### Subspace rotation and dynamics correlates with behavior performance

To test whether the post-training population rotation in the match relative to the nonmatch condition was linked to improved task performance, we compared the subspace rotation angle in correct and error trials in a subset of neurons with sufficient error trials across conditions (n=295 in the spatial working memory task, and n=201 in the feature working memory task), pooling data across all areas (see Methods: Analysis of error trials). In correct trials (Fig. 5A, C), the low dimensional PC subspaces of match and nonmatch epochs were nearly orthogonal to each other (79.9±5.4 and 78.0±14.5 degrees for the spatial and feature stimuli, respectively), consistent with the post-training findings of the full population (Fig. 4). In error trials, this rotation was significantly reduced: Fig. 5B, D,19.7±9.0 and 18.8±14.5 respectively (t-test for spatial: t_100_=57.4, p<0.0001, feature: t_100_=28.9, p<0.0001). To determine how population responses evolved across the length of the trial, we investigated representation dynamics for match and nonmatch trials in the reduced PCA space. Example trajectories for a single stimulus condition for the mid-dorsal area are plotted in Fig. 5E. These also resembled the trajectories in state space described previously for neuronal activity in animals trained to perform perceptual decision tasks ^8^. In correct trials, as expected, match and nonmatch trajectories stayed close to each other until the sample period and then diverged during the sample presentation and the delay period that followed it. In error trials, an abnormal rotation was already present in the cue period and match and nonmatch trajectories were less-distinguishable (Fig. S8). Importantly, in error trials, the neural pattern we observed was not the reverse of that of correct trials. Instead, when the animal wrongly reported the matching status, the match and nonmatch trajectories were similar to each other but distinct from either match or nonmatch condition in correct trials (Fig. S9). The result suggests that incomplete rotation of the stimulus subspace is more likely to result in errors. However, this aberrant trajectory emerged early in the trial and was not specifically dependent on incomplete orthogonalization of the nonmatch stimulus, which, as we showed here, was present even in naïve animals.

**Figure 5.**
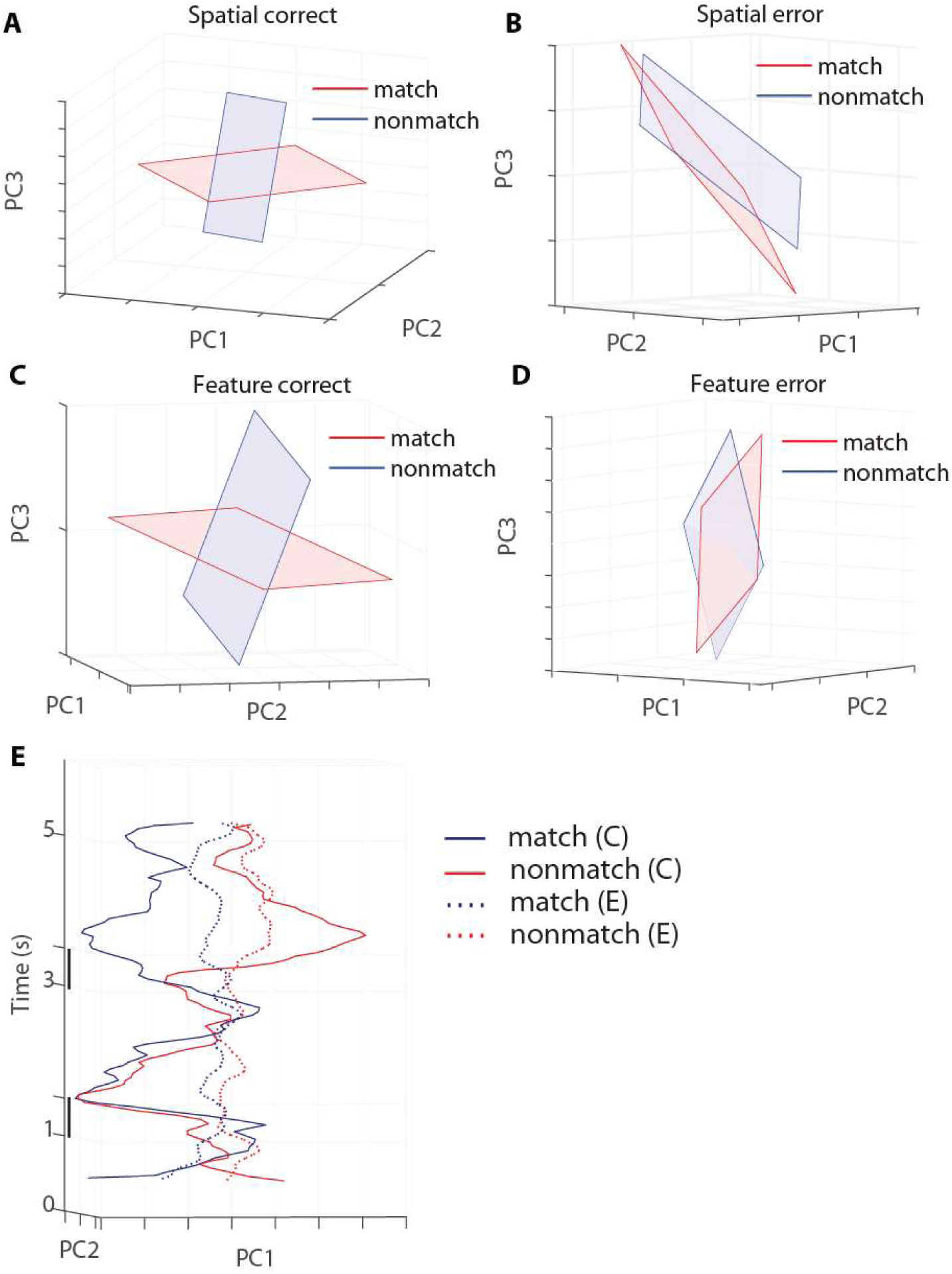
Population representations in correct and error trials. (A) Representation of spatial stimuli during the match and nonmatch period in correct trials (n=295 neurons). Rotation φ=76°. (B) Representation of spatial stimuli during the match and nonmatch period in the error trials (same population as in panel A). Rotation φ=15°. (C) Representation of feature stimuli during the match and nonmatch period in correct trials (n=201 neurons). Rotation φ=84°. (D) Representation of feature stimuli during the match and nonmatch period in error trials (same population as in panel C). Rotation φ=7°. (E) Dynamics of stimulus representations in correct and error trials, for one example spatial location. Solid blue: match in correct trials, solid red: nonmatch in correct trials, dashed blue: match in error trials, dashed red: nonmatch in error trials. The same cells are plotted across all conditions (n=295 neurons).

## DISCUSSION

Computation in neural activity is characterized by the transformation of representations across stages of processing ^12^. Our results demonstrate that neuronal populations transform stimulus representations and exhibit dynamics shown to represent cognitive operations ^5, 8, 9, 10^, even in the absence of task execution, in animals naïve to any such training. Our results in the same dataset from trained animals corroborated previous findings: in some divisions of the PFC, sequential cue and match stimuli from the same trials were represented in orthogonal planes; match and nonmatch stimuli exhibited large state space separations; and error trials exhibited smaller plane rotations than correct ones. Our results indicate that some dynamic transformations of population-based stimulus representations are not caused by task operations requiring explicit training.

### Neural basis of cognitive flexibility

Natural plasticity allows for significant training-induced improvements of working memory, particularly through changes in PFC activity, over long periods of training ^18, 19, 20, 21, 22^. Representation of stimuli in neural activity has also been shown to be dynamic at much faster timescales, during the time course of a single behavioral session ^23^ or, depending on trial events, on a moment-by-moment basis ^1^. The transformation of stimulus representations has been shown to be indicative of flexible task execution, e.g. rotating representations may minimize interference between multiple stimuli, thus allowing additional stimuli to be encoded, or learning of associations between stimuli, otherwise presented passively ^9^. Our current results suggest that the prefrontal cortex may perform these operations automatically, even in the absence of task execution, or learning novel stimuli. Furthermore, the scale of stimulus representation was largely unchanged after training. The idea of stable population information under shifting neural responses over days is an increasingly appreciated phenomenon in studies involving post-learning plasiticity^24, 25^.

Regardless of transformations, data across prefrontal areas and task conditions were generally well fit by low dimensional representations. Low dimensionality in working memory representations is considered a hallmark of generalizability, while high dimensionality is correlated with better discriminability ^26^. In our experiment, stimuli were highly discriminable from each other from the outset, and task training involved incorporating them into the context of a new task rather than learning fine features of the stimuli ^27^, suggesting that an increase in dimensionality was not required to perform the task after training.

### Mechanisms and interpretations of rotations

The mechanisms of the population rotation phenomenon are still under debate. Sequential activation of individual neurons in the motor cortex can create an apparent rotation at the population level ^28^, although the rotation observed experimentally cannot fully be accounted by this phenomenon ^29^. In the prefrontal cortex, neuronal activation is less sequential in nature compared to that of the motor cortex and the orthogonality we observed between contexts could be caused by the activation of largely nonoverlapping coding populations or a change of tuning to stimuli (Fig. S5). Similarly, the interpretation of the functional significance of this low-dimensional rotation, is multifaceted. Previous research has suggested that one of the functions of this rotation may be to reduce the interference between representations, and notably such rotations may appear as mice learn the structure of a stimulus set, without needing to perform an actual comparison based on working memory ^9^. Our finding supports this conclusion by comparing rotations in response to the same stimuli before and after they were incorporated into a working memory tasks. The monkeys in our cohort were familiar with the stimulus sets and structure of the passive “task” (i.e. sequence of two stimulus presentation and their relative timing) by the time recordings in the “naïve” state began. It is therefore possible that the rotations emerged after the animals had discovered this regularity. Future studies may address this question. In any case, our results show that rotations in some PFC regions are task independent and track information related to the temporal structure of the task, by low dimensional rotation, even when that information is not behaviorally relevant. This automatic tracking of variables changing in the natural world without explicitly prompt could be essential for learning.

### Regional Specialization

Prior studies have debated if and how different types of information may be represented across the dorsal-ventral and anterior-posterior axes of the prefrontal cortex has been contentious ^30, 31, 32, 33^. Converging evidence, however, suggests that anterior subdivisions of the prefrontal cortex generally represent more abstract information, and their activation depends to a greater extent on the task subjects have been trained to perform ^34^.

Consistent with this idea, we observed that anterior areas consistently exhibited high rotation values for match and nonmatch stimuli relative to the cue (and relative to each other), both before and after training. Our results raise the possibility that this innate stimulus transformation endows anterior areas of the prefrontal cortex with greater capacity for cognitive flexibility.

## METHODS

Data obtained from six male rhesus monkeys (*Macaca mulatta*), ages 5–9 years old, weighing 5–12 kg, as previously documented ^14^, were analyzed in this study. None of these animals had any prior experimentation experience at the onset of our study. Monkeys were either single-housed or pair-housed in communal rooms with sensory interactions with other monkeys. All experimental procedures followed guidelines set by the U.S. Public Health Service Policy on Humane Care and Use of Laboratory Animals and the National Research Council’s Guide for the Care and Use of Laboratory Animals and were reviewed and approved by the Wake Forest University Institutional Animal Care and Use Committee.

Monkeys sat with their heads fixed in a primate chair while viewing a monitor positioned 68 cm away from their eyes with dim ambient illumination and were required to fixate on a 0.2° white square appearing in the center of the screen. During each trial, the animals were required to maintain fixation while visual stimuli were presented either at a peripheral location or over the fovea, in order to receive a liquid reward (typically fruit juice). Any break of fixation immediately terminated the trial and no reward was given. Eye position was monitored throughout the trial using a non-invasive, infrared eye position scanning system (model RK-716; ISCAN, Burlington, MA). The system achieved a < 0.3° resolution around the center of vision. Eye position was sampled at 240 Hz, digitized and recorded. The visual stimulus display, monitoring of eye position, and synchronization of stimuli with neurophysiological data was performed with in-house software implemented in the MATLAB environment (Mathworks, Natick, MA), utilizing the Psychophysics Toolbox ^35^.

### Pre-training task

Following a brief period of fixation training and acclimation to the stimuli, monkeys were required to fixate on a center position while stimuli were displayed on the screen. The stimuli shown in the pre-training, passive, spatial task consisted of white 2° squares, presented in one of nine possible locations arranged in a 3 × 3 grid with 10° of distance between adjacent stimuli. Only the eight peripheral locations are analyzed here, as the center location never appeared as a nonmatch. The stimuli shown in the pre-training passive feature task consisted of white 2° geometric shapes drawn from a set comprising a circle, a diamond, the letter H, the hashtag symbol, the plus sign, a square, a triangle, and an inverted Y-letter. These stimuli could be presented in one of the possible locations of the spatial grid.

The presentation began with a fixation interval of 1 s where only the fixation point was displayed, followed by a 500 ms stimulus presentation (referred to hereafter as cue), followed by a 1.5 s “delay” epoch where, again, only the fixation point was displayed. A second stimulus (sample) was subsequently shown for 500 ms. In the spatial task, this sample would be either identical in location to the initial stimulus, or diametrically opposite. In the feature task, the sample would appear in the same location as the cue and would either be an identical shape or the corresponding nonmatch shape (each shape was paired with one nonmatch shape). Only one nonmatch sample was paired with each possible cue, so that the number of match and nonmatch trials were balanced in each set. In both the spatial and feature task, this sample stimulus display was followed by another “delay” period of 1.5 s where only the fixation point was displayed. The location and identity of stimuli were of no behavioral relevance to the monkeys during the “pre-training” phase, as fixation was the only necessary action for obtaining rewards.

### Post-training task

Three monkeys, the data of which are analyzed here, were trained to perform active working memory tasks that involved the presentation of identical stimuli as the spatial and feature tasks during the “pre-training” phase. Monkeys were required to remember the spatial location and/or shape of the first presented stimulus, and report whether the second stimulus was identical to the first or not, via saccading to one of two target stimuli (green for matching stimuli, blue for nonmatching). Each target stimulus could appear at one of two locations orthogonal to the cue/sample stimuli, pseudo-randomized in each trial.

### Surgery and neurophysiology

A 20 mm diameter craniotomy was performed over the PFC and a recording cylinder was implanted over the site. The location of the cylinder was visualized through anatomical magnetic resonance imaging (MRI) and stereotaxic coordinates post-surgery. For two of the four monkeys that were trained to complete active spatial and feature working memory tasks, the recording cylinder was moved after an initial round of recordings in the post-training phase to sample an additional surface of the PFC.

### Anatomical localization

Each monkey underwent an MRI scan prior to neurophysiological recordings. Electrode penetrations were mapped onto the cortical surface. We identified six lateral PFC regions: a posterior-dorsal region that included area 8A, a mid-dorsal region that included areas 8B and 9/ 46, an anterior-dorsal region that included area 9 and area 46, a posterior-ventral region that included area 45, an anterior-ventral region that included area 47/12, and a frontopolar region that included area 10. However, the frontopolar region was not sampled sufficiently to be included in the present analyses.

### Neuronal recordings

Neural recordings were carried out in the aforementioned areas of the PFC both before and after training in each WM task. Extracellular recordings were performed with multiple microelectrodes that were either glass- or epoxylite-coated tungsten, with a 250 μm diameter and 1–4 MΩ impedance at 1 kHz (Alpha-Omega Engineering, Nazareth, Israel). A Microdrive system (EPS drive, Alpha-Omega Engineering) advanced arrays of up to 8 microelectrodes, spaced 0.2–1.5 mm apart, through the dura and into the PFC. The signal from each electrode was amplified and band-pass filtered between 500 Hz and 8 kHz while being recorded with a modular data acquisition system (APM system, FHC, Bowdoin, ME). Waveforms that exceeded a user-defined threshold were sampled at 25 μs resolution, digitized, and stored for offline analysis. Neurons were sampled in an unbiased fashion, collecting data from all units isolated from our electrodes, with no regard to the response properties of the isolated neurons. A semi-automated cluster analysis relied on the KlustaKwik algorithm, which applied principal component analysis of the waveforms to sort recorded spike waveforms into separate units. To ensure a stable firing rate in the analyzed recordings, we identified recordings in which a significant effect of trial sequence was evident at the baseline firing rate (ANOVA, p < 0.05), e.g., due to a neuron disappearing or appearing during a run, as we were collecting data from multiple electrodes. Data from these sessions were truncated so that analysis was only performed on a range of trials with stable firing rates. Less than 10% of neurons were corrected in this way. Identical data collection procedures, recording equipment, and spike sorting algorithms were used before and after training in order to prevent any analytical confounds.

### Dimensionality reduction and rotation of subspaces

We applied principal components analysis (PCA) to visualize the neural population activity manifolds during the spatial and feature tasks. PCA was performed on the mean firing rate of neurons across different prefrontal regions, for data collected both before and after training. We examined the submanifolds of neural activity as a function of the spatial and feature stimulus sets.

In the spatial task, we collected data from five prefrontal regions before and after the monkeys were trained to perform the task. In each region, trials were collected when the cue stimuli appeared at *L* = 8 locations. We then selected the recordings with more than 16 trials at each location. For each neuron, the average firing rate of the match trials (and nonmatch trials) during a given period at the eight locations formed a column entry (8×1 vector) for the population activity matrix *A*_l_ (*A*_2_ for nonmatch trials). During each task epoch we defined the corresponding activity matrix *A*_l_ (*A*_2_) for each prefrontal region as an 8 × *N* matrix, where 8 is the number of locations where the cue occurred and *N* is the number of neurons. To find the rotation angle between low-dimensional representations in two task epochs, we aligned the activity matrix into one single matrix *B*, which is a 16 × *N* matrix with the first eight rows containing the activities in one period and the rest containing another period. Then we normalized *B* by subtracting the mean across each column to guarantee the matrix was zero-centered. PCA was applied to the centered data using singular value decomposition (SVD). We selected the first three eigenvectors of the covariance matrix of the zero-centered data and sorted them by decreasing order in terms of explaining the variance. The first three principal components explained an average of 73% of the response matrix variance across all examined periods and locations. Similar procedures were applied to visualize the responses in the feature task, where we grouped the data according to the shape of the cue stimulus. Specifically, there were eight different shapes in the feature task. Therefore, the corresponding activity matrix had a dimension of 8 × *N*, in which *N* is the number of neurons, and each row stands for a shape in the feature task (see Fig. 1B). For each pair of comparison for the rotations, we aligned the data in the same way as described above.

To visualize population representations, we projected the centered activity matrix into a three-dimensional (3D) PCA space. After projection, the population representations of the eight spatial locations (and eight shapes in the feature task) formed eight points conjugated on a 2D plane in the reduced 3D space. Following a similar procedure, we further applied PCA on the eight points in the reduced plane, with a dimension of 8 × 3, to find its best-fit plane. The high proportion of variance explained by the fitted plane (e.g. for the PD area in the cue presentation epoch pre-training: 89.1%±3.3%; post-training PD 85.1%±3.6%) validated our hypothesis that the reduced space could be presented in 2D.

We defined the angles between different conditions based on the best-fit planes that we constructed above. Given the vectors spanning the best-fit plane (*P*_l_) for one period are 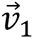 and 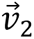 (obtained from the PCs of the reduced eight points), and the best-fit plane (*P*_2_) for another period in the same region are 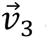 and 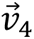, the angle between *P*_l_ and *P*_2_ was calculated as follows:

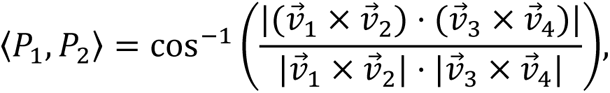

where “〈*P*_l_, *P*_2_〉” denotes the angle between *P*_l_ and *P*_2_, cos^-l^ is the inverse cosine function, and 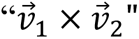 is the cross product that finds the vector that is perpendicular to the plane spanned by 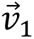 and 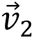, the “⋅” sign stands for the dot product, and “|x|” returns the absolute value (for a scaler) or length (for a vector) of x.

To distinguish between an authentic large rotation of stimulus representation and an incidental rotation arising in a population of neurons with low selectivity and high variability, we employed a method based on synthetic data to create a control condition for each rotation-angle measurement. Specifically, we calculated the mean firing rate from n trials for each cell and each stimulus class across different task epochs. For example, we computed the mean firing rate during the cue period from the 16 trials of cell #1 located in the anterior dorsal area during the pre-training spatial task and stored this mean as *λ*. Subsequently, we generated a Poisson distribution with the same mean (*λ*) and randomly produced n sample values from it, simulating n trials from the cue period of this specific cell#1 in the anterior dorsal area during the pre-training spatial task. This control dataset mirrored the variability and selectivity statistics of the empirical dataset, thus forming a baseline for rotation within each epoch, and providing a reference for angle and geometry measurements. Control conditions for rotation angle and geometry measurements are plotted with dashed boxes, whereas empirical data are indicated by solid frame boxes, as illustrated in Fig. 3B1-B2 and Fig. 4. Only cells with at least 6 match and 6 nonmatch trials for each stimulus condition were included in the rotation angle analysis. We assessed variability by randomly drawing 80% of the cells from each prefrontal subdivision over 1000 iterations.

We implemented a nonparametric bootstrap method to assess the statistical significance of the differences between empirical angles. First, the underlying true angle difference was calculated using the angle difference derived from all trials and cells.

Then we conducted the bootstrap procedure. In each iteration, we shuffled the training statuses and subdivision labels of all cells under comparison, followed by the recalculation the angle difference post-shuffle. This process was repeated 1000 times. The p-value for the observed difference was estimated based on the proportion of values in the shuffled data, that exceeded true empirical value.

### Variance accounted for ratio (VAF)

To cross-check the rotation angle between the 2D subspace of different task epochs, we measured the Variance Accounted For (VAF) ratio for each angle measurement^15^. The VAF ratio for epoch subspace pair (1,2) was defined as follows:

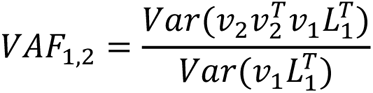

Where *ν*_i_ (*i* = 1,2) is a 3×2 matrix representing two subspace axes, and *L*_l_ represents the stimulus projection in the first subspace. VAF ranges from 0 to 1; large values indicate better alignment of subspaces, while values close to 0 suggest orthogonality. Only cells with at least 6 match and 6 nonmatch trials for each stimulus category were included in the VAF analysis.

### Single cell basis of coding subspace

To characterize the geometric relationship between single-cell axes and the low-dimensional subspace in certain contexts, we aimed to measure the alignment of single cells’ selectivity with the low dimensional subspace. Geometrically, better alignment corresponds to a smaller angle between a vector and a subspace. This also implies that variance in the vector would yield significant variance within the subspace. Intuitively, a better-aligned vector will have a larger projection onto the subspace.^15^. In accordance with this concept, we projected a unit vector *i* from the neuron under question onto the coding subspace of a specific context. The resultant projection vector onto the subspace is denoted by *A*, and the angle between *A* and a specific stimulus vector (we chose the first location/shape) is *φ*, so *A* and *φ* satisfy:

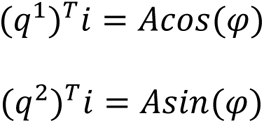

in which *q*^1^ and *q*^2^ represent the first two PCs of the subspace. Thus *A* measures the degree of alignment, or the strength of contribution from the cell, and *φ* indicates the turning direction of the cell in the coding subspace. We also employed a quantity called normalized participation ratio (PR) ^15^ to quantify how distributed the subspace is across the population, as follows:

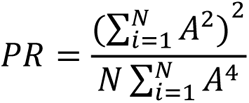

This quantity ranges from 0 to 1, with 0 indicating very sparse coding and 1 indicating evenly distributed coding across the whole population.

To examine the contribution of repetition suppression on the observed subspace rotation, cells with prominent repetition suppression were removed from the database, and the rotation angle was recalculated. Specifically, the top 10% of cells with the largest reduction in firing rate from the cue to the match epoch was excluded from this analysis.

### Geometrical order in low-dimensional representation

A matrix *L* of size 8 × 2 was used to represent the projection of 8 locations or shapes on the 2D plane, with each row representing a stimulus in the 2D PCA space. To compare geometric order with a hypothetical order on an octagon (Fig. 2, Fig. S10), we first projected the 8 stimuli onto a circle. Specifically, we aligned the 8 points uniformly on an imaginary circle by equalizing the arrangement as follows

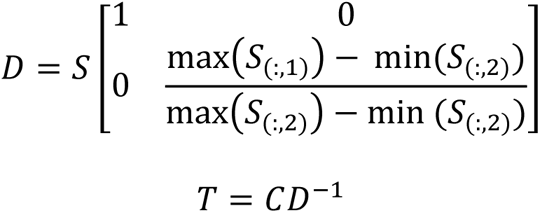

Here, S represents the score matrix and C the coefficient matrix, both obtained from the PCA of L. Here, S_(:,i)_ is the *i*th column of S, and T is the new matrix representing the 8 stimuli in the reduced space. Subsequently, we determined the circular order of the stimuli by computing the angles of vectors from the center to each point, with the angle for the *i*th stimulus defined as θ_i_=arccot(*I*), where *I* denotes the vector from the center to the *i*th point.

Focusing solely on the sequential order, we aligned the stimuli to the vertices of an ideal octagon in the order obtained from the circular projection. This realignment is defined as N(o_rank_,:)=M(o_hypothetical_,:), where o_rank_ and o_hypothetical_ are vectors containing the measured stimuli order in the low dimensional space and the hypothetical order, respectively. M contains coordinates of a unit-size octagon, and N is a low-dimensional representation of the 8 stimuli, reshaped to a standard octagon.

We quantified "order regularity" by the unexplained variance from the Kabsch algorithm ^36^, which finds the optimal rotation matrix R that minimizes the root mean squared deviation between M and N. The Kabsch algorithm was implemented as follows

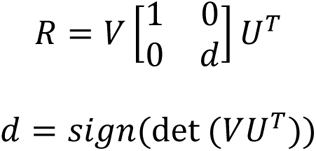

where U and V come from the singular value decomposition of the covariance matrix of M and N. Unexplained variance between two set of coordinates, referred to as C_first_ and C_second,_ is defined as

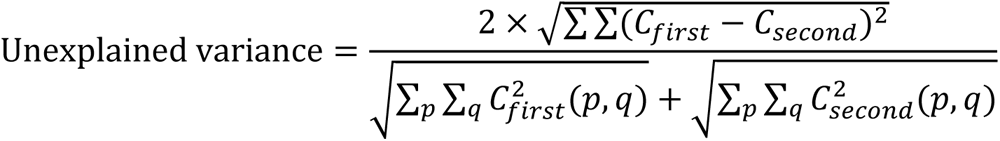

A lower value of unexplained variance indicates a higher congruence, signifying greater "order regularity."

When comparing the empirical geometry with a hypothetical geometry (Fig. 2, Fig. S10), we hypothesized that the distance between each cue and its nonmatch pair should be maximized. For the spatial task, the match-nonmatch structure is symmetric (e.g. when A is cue, B is nonmatch; and when B is cue A is nonmatch). These two assumptions should lead to a predicted arrangement similar to the physical appearance of the spatial stimuli, where the nonmatch pair always perfectly falls on two ends of a diagonal line.

Only cells with at least 6 match and 6 nonmatch trials for each stimulus category were included in the geometrical similarity analysis. Standard error was measured by random drawing 80% of cells in each subdivision over 100 iterations.

### Time course of subspace rotation

In addition to visualizing the rotation between different phases of the WM task, we investigated the dynamics of presentations in the reduced PCA space. To construct the trajectory, we discretized the whole task with a timestep of Δ*_t_*= 50 *ms* and an interval of *l_t_* = 250 *ms*. For each timestep k, we recorded the mean firing rates for the match and nonmatch trials in the interval [*t*_0_ + (*k* − 1)Δ*_t_* − 250, *t*_0_ + (*k* − 1)Δ*_t_*] as the entries for the population activity matrix *A*_l_ and *A*_2_ (defined in the above section). Similar as the procedure of constructing the PCA space, we projected the neural responses for each condition into their first three principal components. To plot the trajectories, we started from 250 milliseconds after the start of fixation, i.e., set the initial time *t*_0_ = 250 *ms*. Consequently, the first interval we considered spans from 0 to 250 milliseconds in fixation. We also used the first three PCAs in the match trials during the cue period as the common basis to calculate the coordinates. In other words, we projected the activity matrix into the same common subspace spanned by the PCA space of the match trials. As illustrated in Fig. 3A2, we explored the dynamics of representation for each location (shape) in the spatial (feature) task, where we plot every six points from the dynamics, i.e., we set an increment of *dt* = 300 *ms* for the purpose of visualization.

To further investigate the changes in decoding spaces, we randomly sampled half of the trials from all periods (fixation, cue, delay1, sample and delay2) and calculated its PCA space as the base to project the remaining half trials. The area of the projection for a given period indicates how much information the state space contains from all eight locations, which we defined as the decoding subspace (S_t_). For example, in the reduced space formed by PC1 and PC2, the eight spatial task locations were presented by eight points. To quantify the size of our decoding subspace, we first found the boundaries of the eight points, which was a set of points representing a single conforming to a 2-D boundary around the eight points. Then we calculated the areas of the polygons defined by the boundary points, which we defined as the areas of the decoding space (A_t_). We calculated the ratios of the areas of decoding spaces based on the fixation period (A*_fix_*). For example, the ratios (*R*) between fixation, delay1 and delay2 were defined as follows:

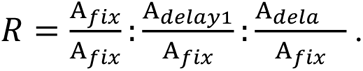

Note that we divided each area by that computed at the fixation period to normalize the data so that we could compare the ratios from different samples. We repeated the random sampling across 10,000 iterations, and the statistics of the ratios were reported in the results section.

### Significance testing of rotations with bootstrap

Our previous analysis was based on all data recorded from neurons that had more than 16 trials for all cue conditions and in each region in Fig. 1C. To test the significance and robustness of the representation rotations, we applied a bootstrap method to sample 100 iterations from the original data. On each iteration, 80% of the cells were drawn with replacement. The angles were calculated following the same procedure as above. Mean and median angles from 100 iterations obtained from bootstrap were represented by dots and horizontal bars within the box plots, while the interquartile range and five percent extreme values were indicated by box border and whiskers, respectively.

### Analysis of error trials

In the post-training spatial task, most cells did not have an error trial. Therefore, the corresponding activity matrix (as defined above) for the error trials was very sparse. In the spatial task, there were 295 neurons where error trials presented at more than half of the eight locations, i.e., 295 rows of the population activity matrix had more than four non-empty entries. We selected those 295 neurons for the error trial analysis in the spatial task. Following the same criterion, we selected 201 neurons containing error trials in the feature task. We first aligned the activity matrix for both cases in the same order, i.e., the same neuron occupied the same row in both matrices. In most cases, for a given stimulus, there were more correct trials than error trials, which led to a larger number of samples in most entries of the activity matrix. Each entry of the activity matrix was calculated from the mean activity across all trials. We thus matched the number of trials in each entry to allow comparison between correct and error trials. Specifically, for each empty entry in the error matrix, we removed the corresponding entry in the correct matrix. Additionally, we selected the same number (minimum of the corresponding entries) of trials in both cases to ensure the same number of samples in both the correct and error trials.

Dimensionality reduction was performed on the error trials using PCA. Given that there are several possible versions of PCA for a sparse matrix, we relied on a probabilistic principal component (PPCA) with variational Bayesian learning ^37^ as this method worked best for our neural data.

## Acknowledgments

Research reported in this paper was supported by the National Eye Institute of the National Institutes of Health under award number R01 EY017077. We wish to thank Rye Jaffe, Junda Zhu, and Zhengyang Wang for helpful comments on the manuscript.

## Author Contributions

C.C. conceived and designed the experiments. C.C. and X. L. Q. performed experiments. S.P, W.D., and C.C. performed data analysis. C.C., S. P. and W.D. wrote the manuscript with input from all authors.

## Declaration of Interests

The authors declare no competing interest.

## Supplementary Information

<u>Supplementary Figures S1-S10</u>

<u>Supplementary Tables S1-S5</u>

**Supplementary Figure S1.**
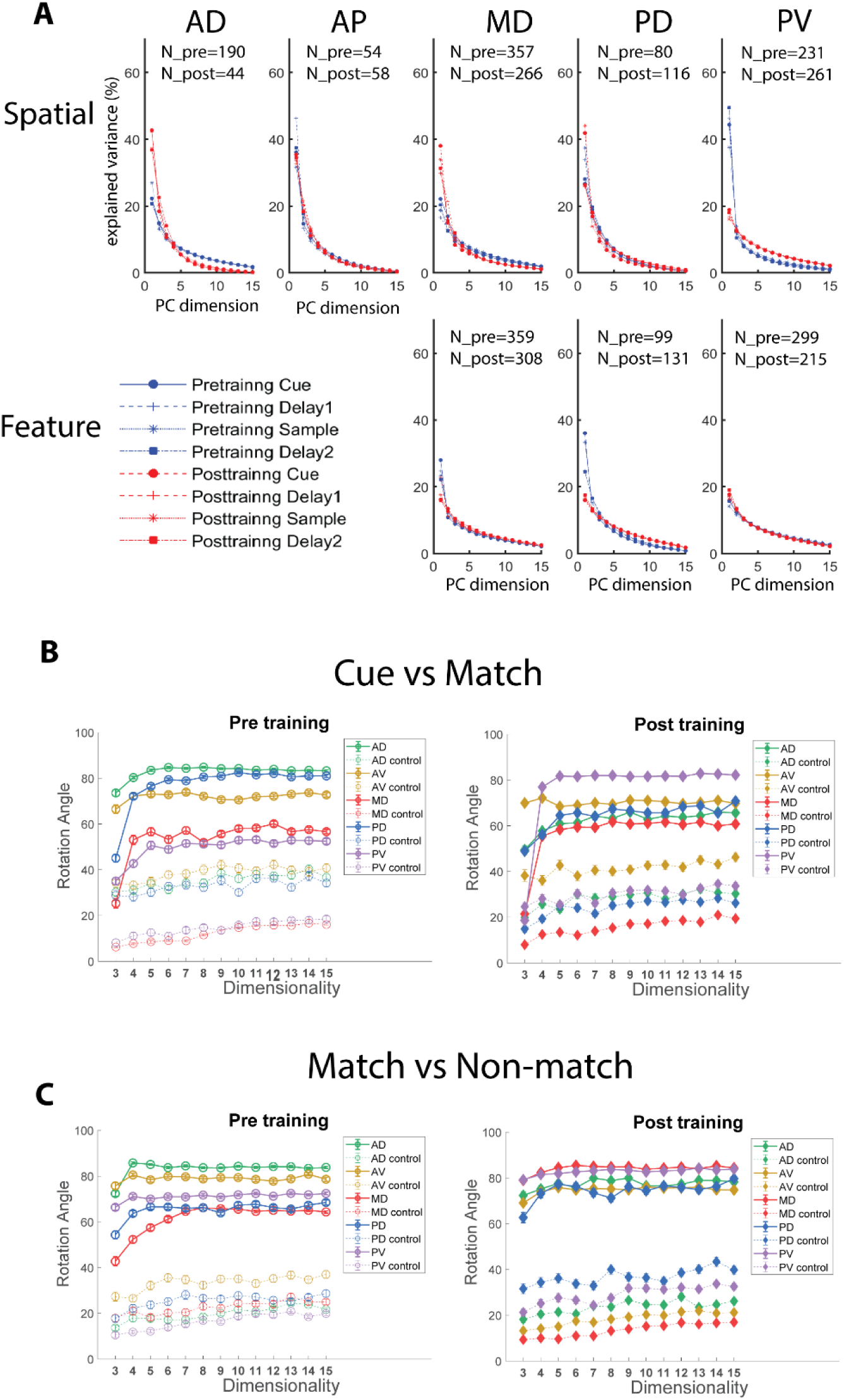
Explained variance and rotation angles in higher dimensions. (A) Explained variance distribution for both tasks pre- and post-training in the PCA analysis. In general, fewer dimensions were needed for characterizing population activity in the spatial task, but under most conditions, three dimensions were enough to explain more than 50% of the variance. The training process has a heterogenous effect on the distribution of variance in different subdivisions of LFPC. (B, C) Angle measurements between 2d planes in 3-15 dimensional space, for cue vs. match and match vs. nonmatch comparison in the spatial dataset. Due to the nature of high dimensional spaces, angle measurements in those spaces tend to be closer to orthogonal, shown by the increasing trend of the curves. Nonetheless, results in low and high dimension spaces are qualitatively similar, indicated by parallel between the curves in most cases.

**Supplementary Figure S2.**
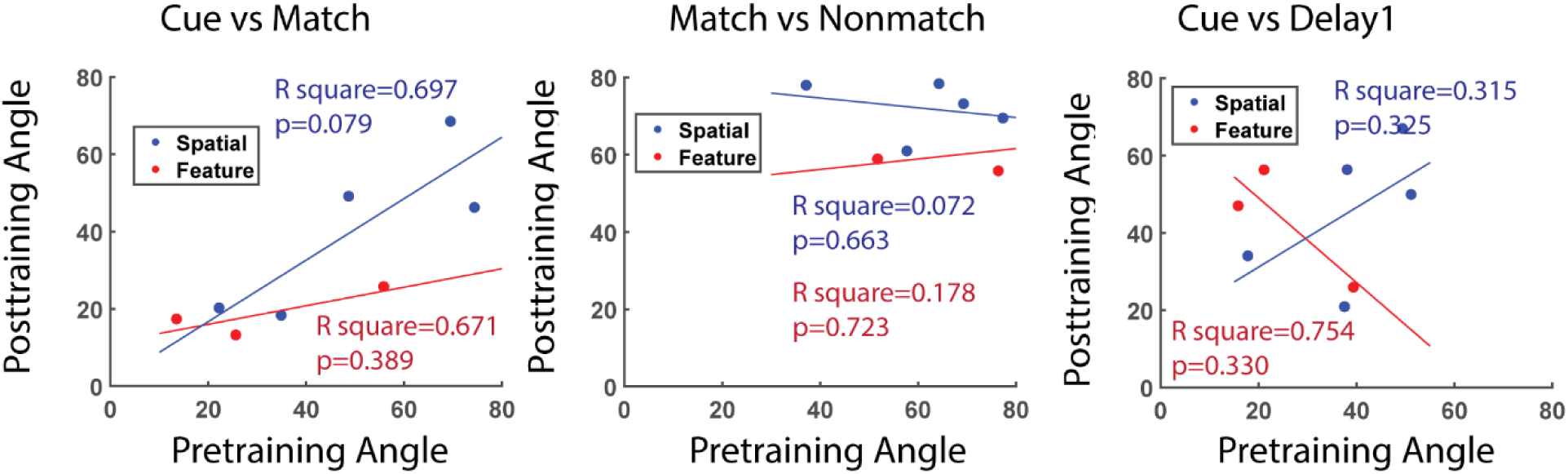
Rotation angles across prefrontal areas are correlated before and after training. Scatter plot of subspace rotation angle for the spatial and the feature task. Each dot represents the mean angle measure for one PFC subdivision, blue color dots indicate spatial task, and red dots indicate feature task.

**Supplementary Figure S3.**
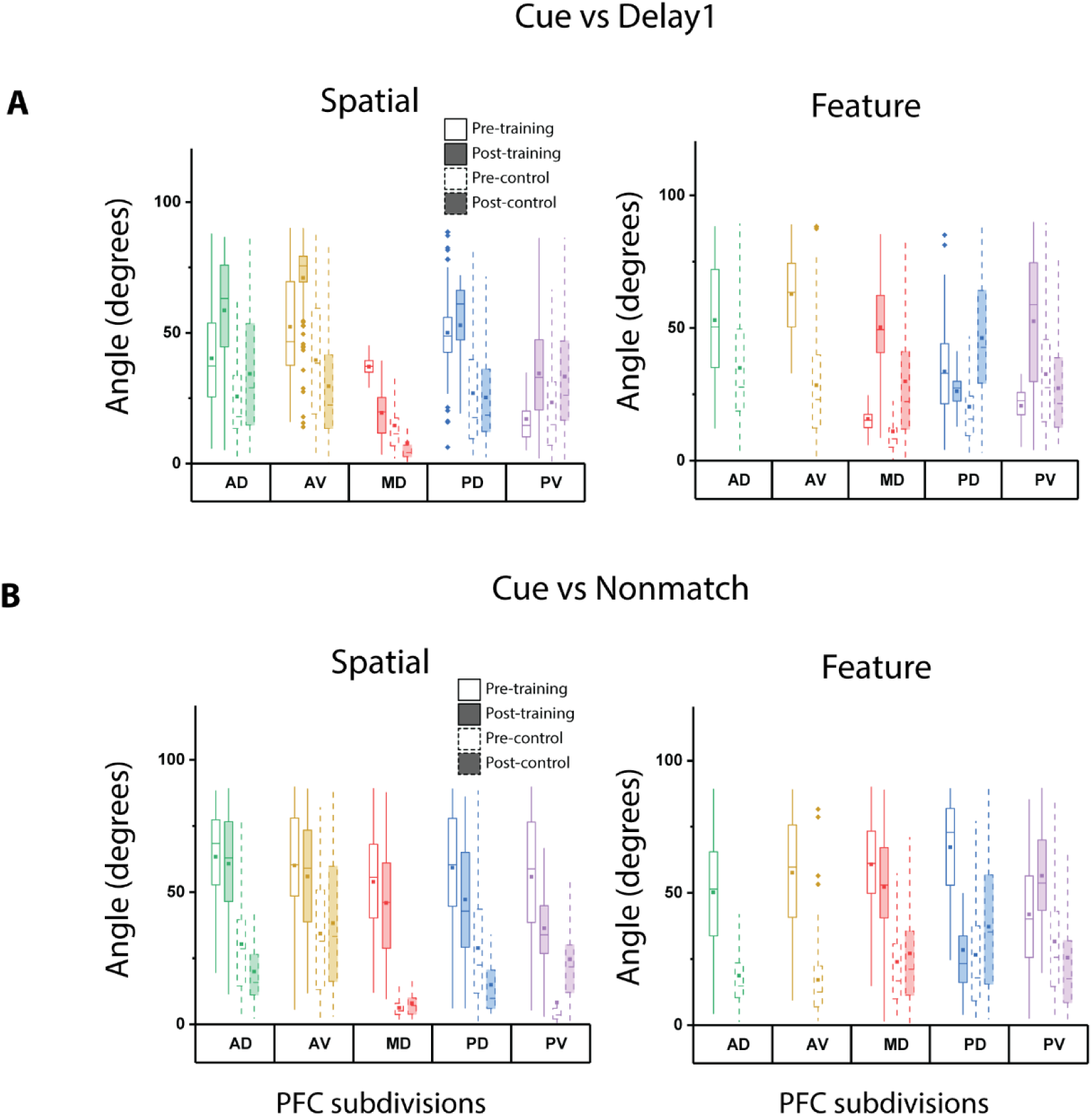
Subspace rotations and geometrical similarity comparisons for different context pairs. (A) Subspace rotation between the cue and the first delay. Plot conventions are the same as Fig. 2-4. (C) Subspace rotation between the cue and the nonmatch trials in the sample period. Plot conventions are the same as Fig. 2-4.

**Supplementary Figure S4.**
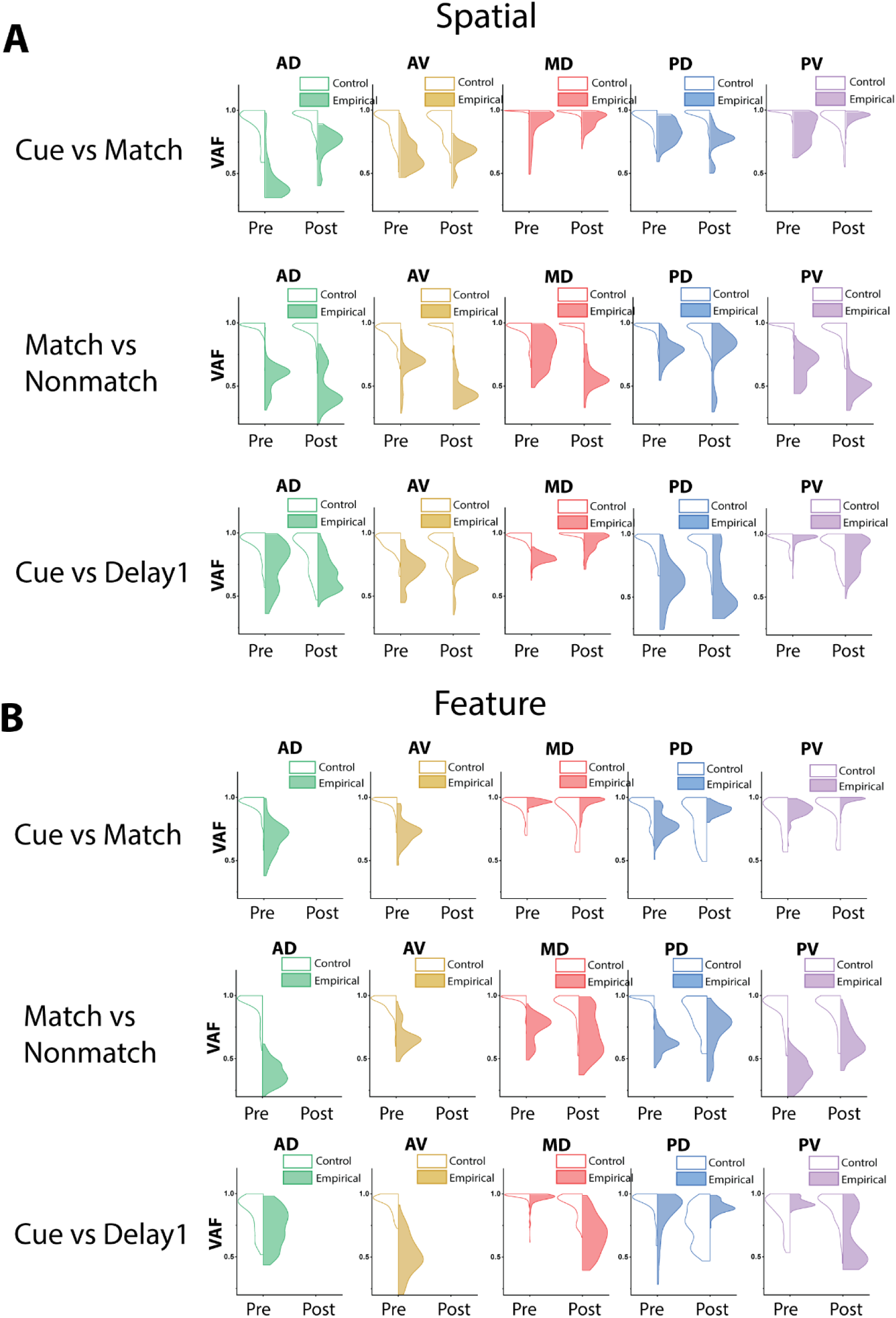
Variance Accounted For ratio distributions across different subdivision for various context comparisons. Variance accounted for (VAF) ratio between different context pairs (top: cue vs. match; mid: match vs nonmatch; bottom: cue vs delay1) in the spatial and the feature task. Compare with angle measurements in Fig. 3, 4, S3. Color filled distributions represent empirical data, while empty distributions represent control measurements where the measure was taken within the same task epoch.

**Supplementary Figure S5.**
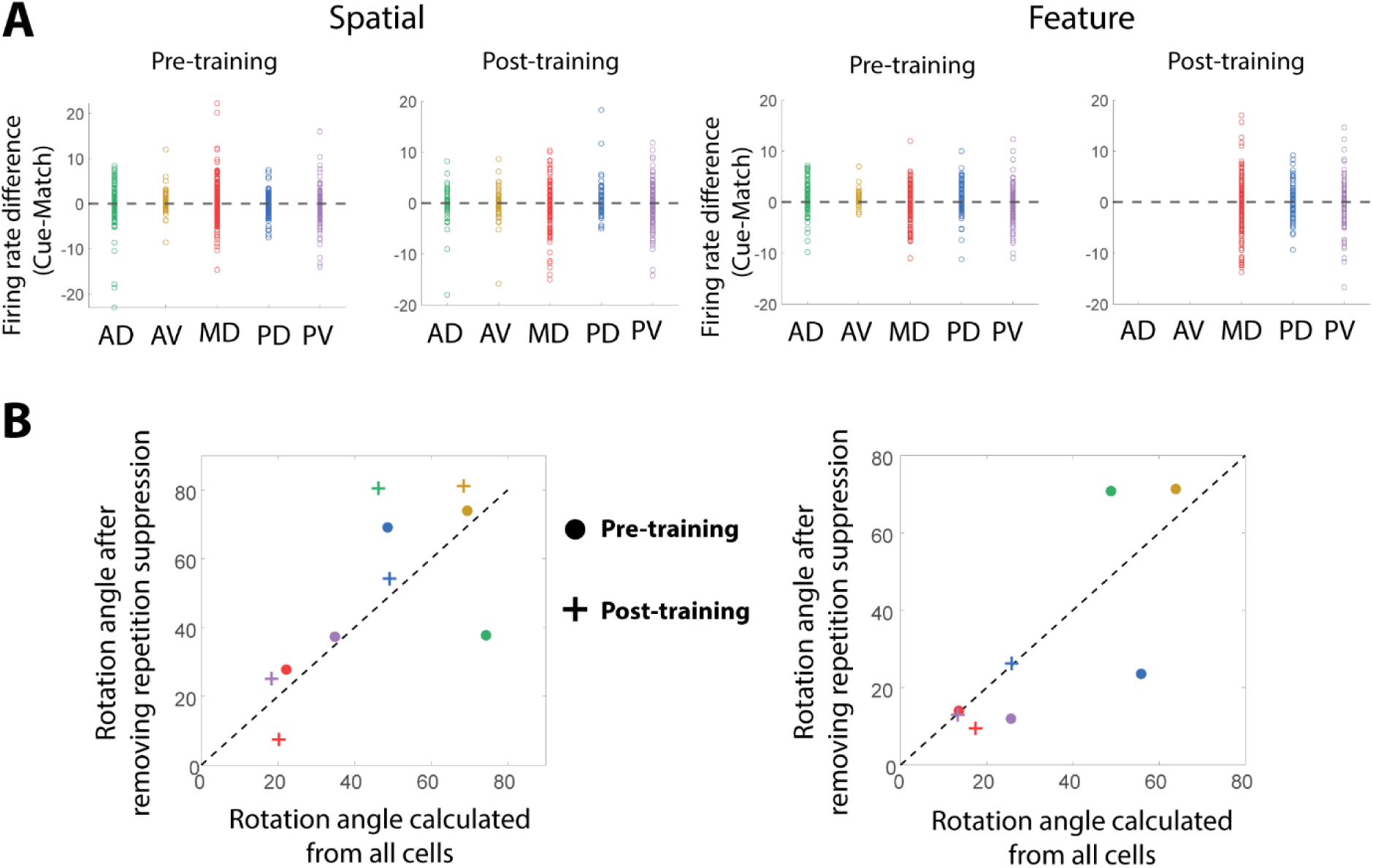
Observed rotations were not caused by repetition effect of single cells. (A) Raw firing rate difference between the cue and match period for all cells in different subregions of PFC. No strong repetition effect was observed, indicated by zeros mean for firing rate differences in all areas. (B) After removing 10% cells with the largest firing rate differences between the cue and the match period, angle measurements are still correlated with the original dataset. Color codes for both (A) and (B) are the same as in Fig. 3B1-B2.

**Supplementary Figure S6.**
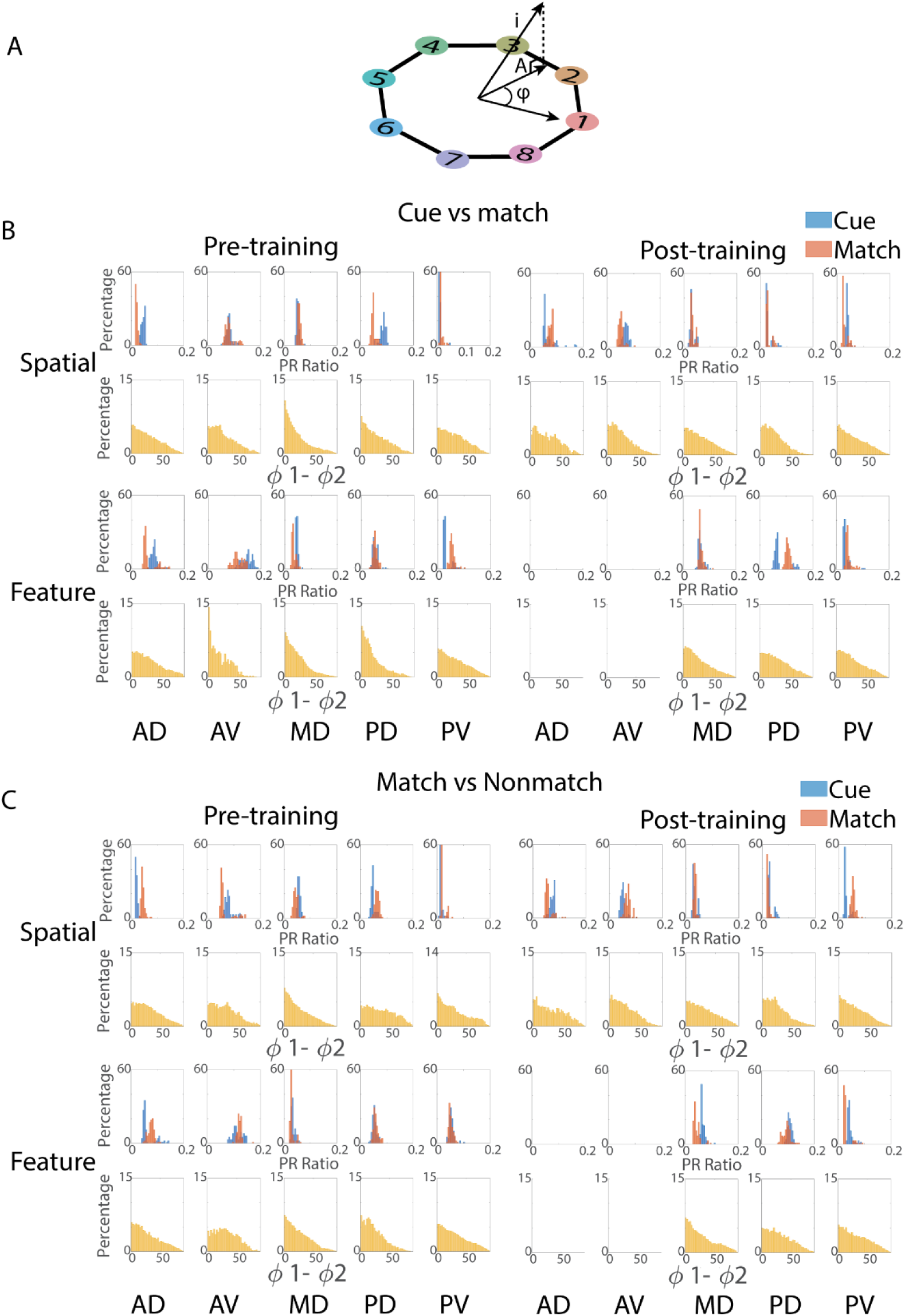
Single cell’s alignment to subspace. (A) Diagram of single cell alignment to a subspace. A unit-length vector representing single cell response was projected back to a specific subspace. Projection A in the subspace represents the degree of alignment to the subspace, while selectivity was quantified by the angle (φ) between the projection vector and certain stimuli in the space. (B-C) Participation ratio (PR) and selectivity change analysis for two contexts comparisons (cue vs. match & match vs. nonmatch) in the spatial and feature task. The PR quantified the proportion of the population that participated in subspace coding. PR values close to 1 indicate more evenly distributed coding across the population, while a PR measurement close to 0 indicates sparse coding. The φ difference (φ1-φ2) between two contexts (cue vs. match & match vs. nonmatch), represents the stimulus tuning changes in different contexts.

**Supplementary Figure S7.**
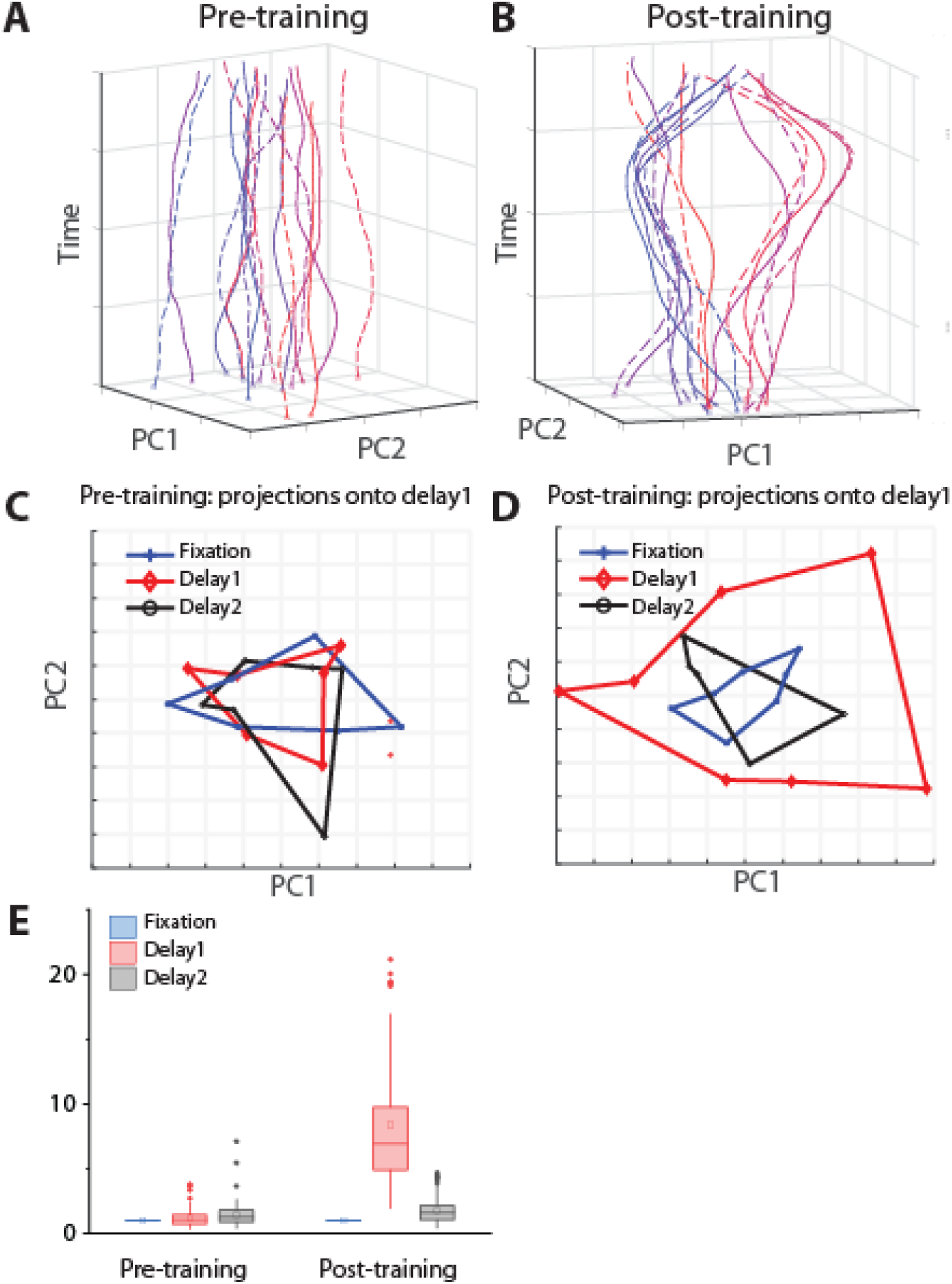
Complete population activity trajectories before and after training. (A) Trajectory of population representation in state space prior to training, for the spatial stimuli in the mid dorsal area (n=453). Solid lines: match trials, dashed lines: nonmatch trials. (B) As in A, for neuronal activity in the mid-dorsal area after training (n=374). (C) Area covered by projection of spatial stimuli prior to training, in three task epochs: fixation, delay1, and delay2. (D) As in C, for data obtained after training. (E). Boxplot of surface areas: summarizing statistics of panels C and D with bootstrapping sampling, where each sample contained 80% of cells. Area values have been normalized by that of the fixation period.

**Supplementary Figure S8.**
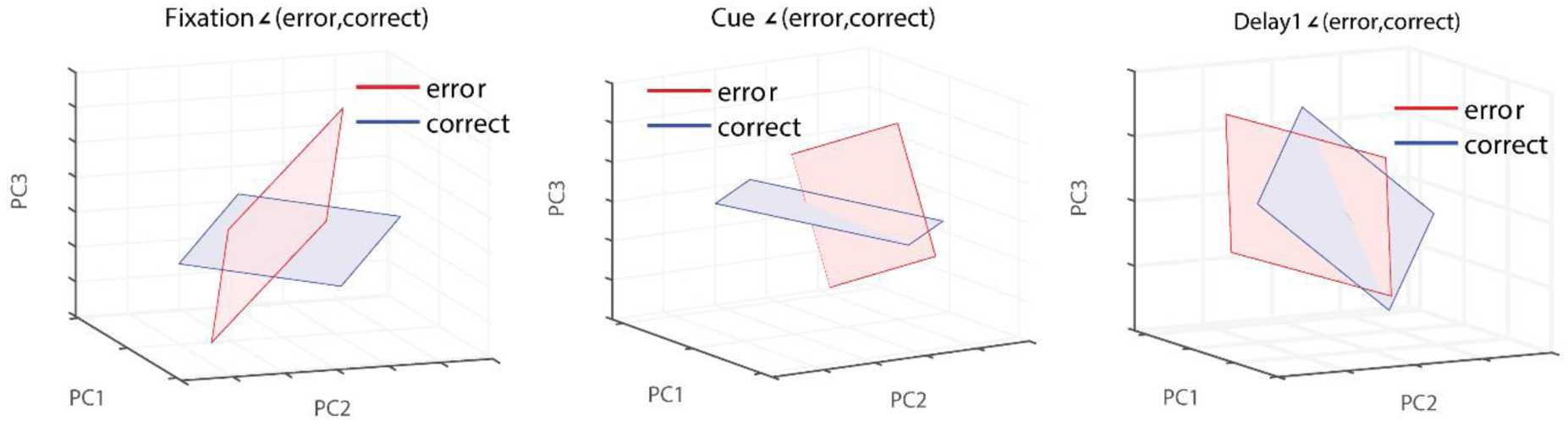
Population subspace rotation in correct and error trials. (A) Representation of spatial stimuli during the fixation period in correct and error trials (n=295 neurons). Rotation φ=45°. (B). As in panel A, for the cue period. Rotation φ=81°. (C) As in panel A for the first delay period. Rotation φ=82°.

**Supplementary Figure S9.**
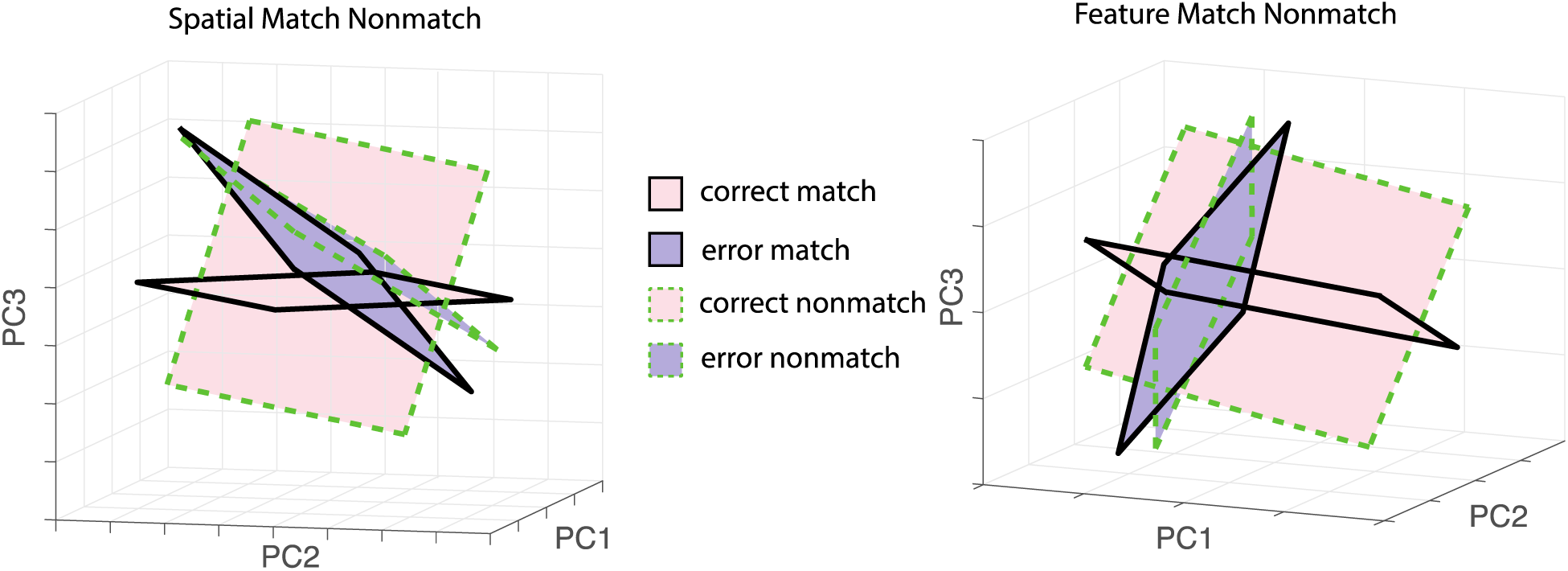
**Population subspace rotation** of stimuli during the match and non-match trial in correct and error trials for the spatial (left) and the feature (right) task.

**Supplementary Figure S10.**
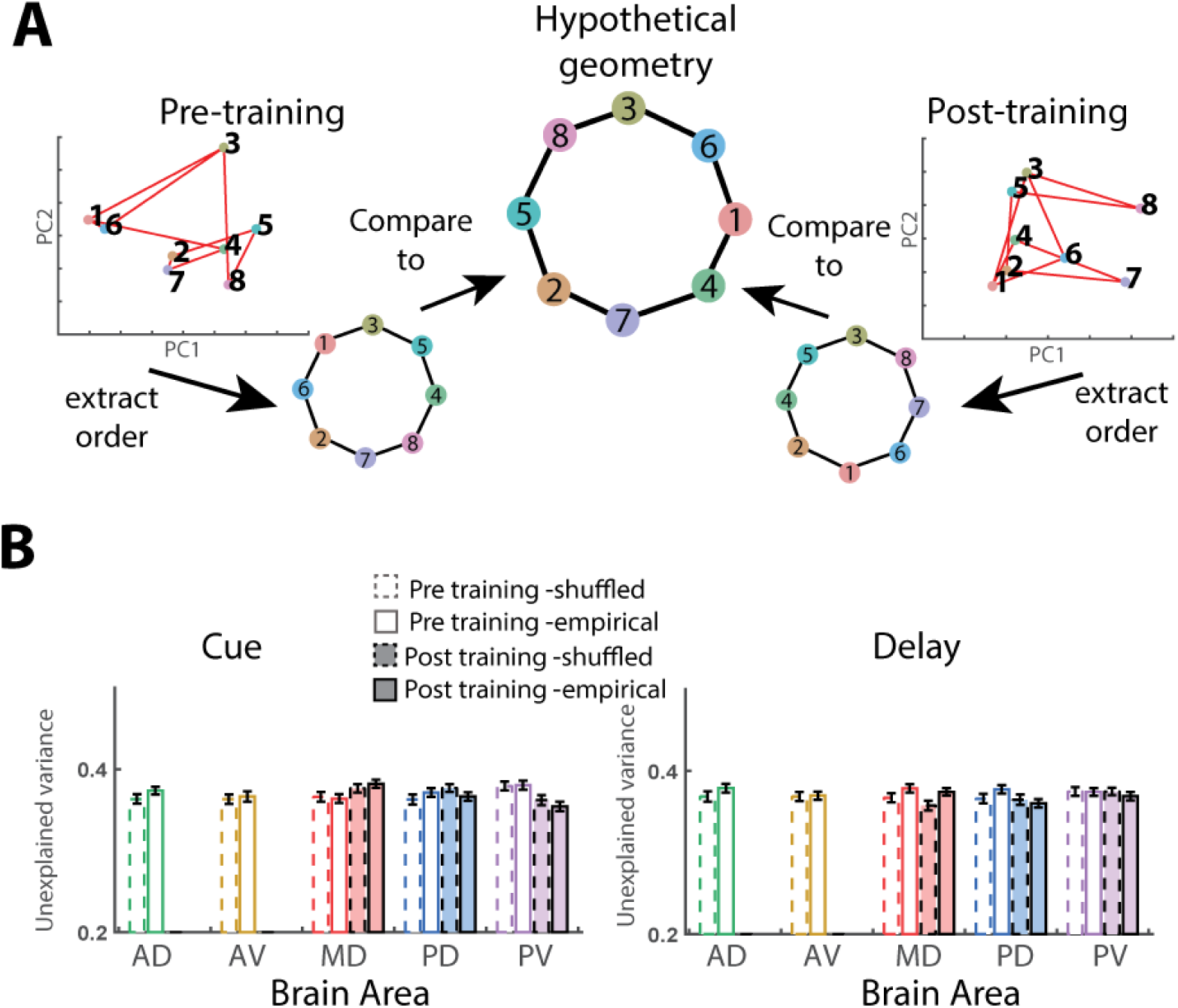
Unexplained variance measurements for feature task in the cue and the delay period, plotting convention the same as Fig. 2. In both task epochs there no area shows significant order regularity based on the match-nonmatch stimuli pairing.

**Table S1:**
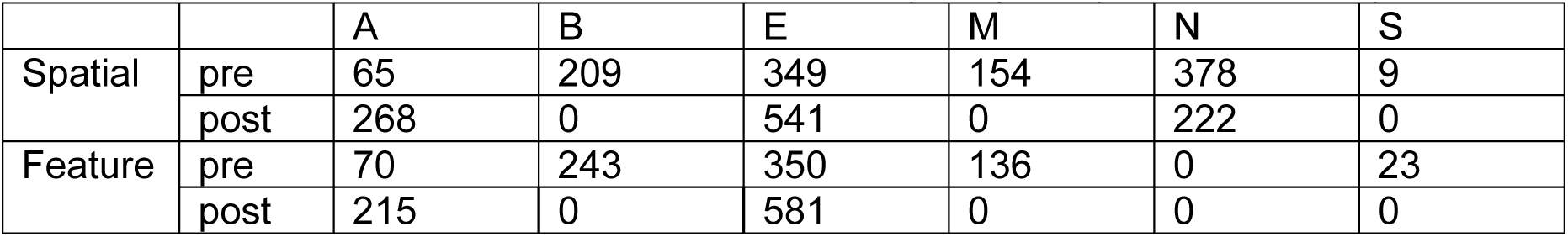
Neuron sample sizes from different monkey subjects (A, B, E, M, N, S)

|  |  | A | B | E | M | N | S |
| --- | --- | --- | --- | --- | --- | --- | --- |
| Spatial | pre | 65 | 209 | 349 | 154 | 378 | 9 |
|  | post | 268 | 0 | 541 | 0 | 222 | 0 |
| Feature | pre | 70 | 243 | 350 | 136 | 0 | 23 |
|  | post | 215 | 0 | 581 | 0 | 0 | 0 |

**Table S2:** Variance explained by the 2-D fitted plane in three dimensional space.

|  |  |  | AD | AV | MD | PD | PV |
| --- | --- | --- | --- | --- | --- | --- | --- |
| Pre-training | Spatial | Cue | 73.3%±1.8% | 78.6%±3.2% | 78.1%±1.6% | 89.1%±3.3% | 80.7%±3.3% |
|  |  | Match | 81.1%±1.4% | 81.5%±2.2% | 74.2%±1.5% | 83.0%±2.3% | 82.8%±3.3% |
|  | Feature | Cue | 76.3%±2.0% | 83.9%±4.2% | 79.9%±1.6% | 79.9%±3.6% | 73.4%±1.4% |
|  |  | Match | 78.4%±2.4% | 80.2%±4.1% | 78.3%±1.7% | 82.4%±2.5% | 73.7%±1.3% |
| Post-training | Spatial | Cue | 80.7%±3.3% | 78.7%±2.9% | 84.0%±1.4% | 85.1%±3.6% | 73.6%±1.4% |
|  |  | Match | 83.6%±1.8% | 86.2%±2.2% | 79.4%±1.9% | 84.2%±3.9% | 73.5%±2.5% |
|  | Feature | Cue |  |  | 72.4%±1.5% | 73.7%±1.6% | 75.5%±1.8% |
|  |  | Match |  |  | 72.9%±1.3% | 75.1%±1.5% | 73.8%±1.7% |

**Table S3:**
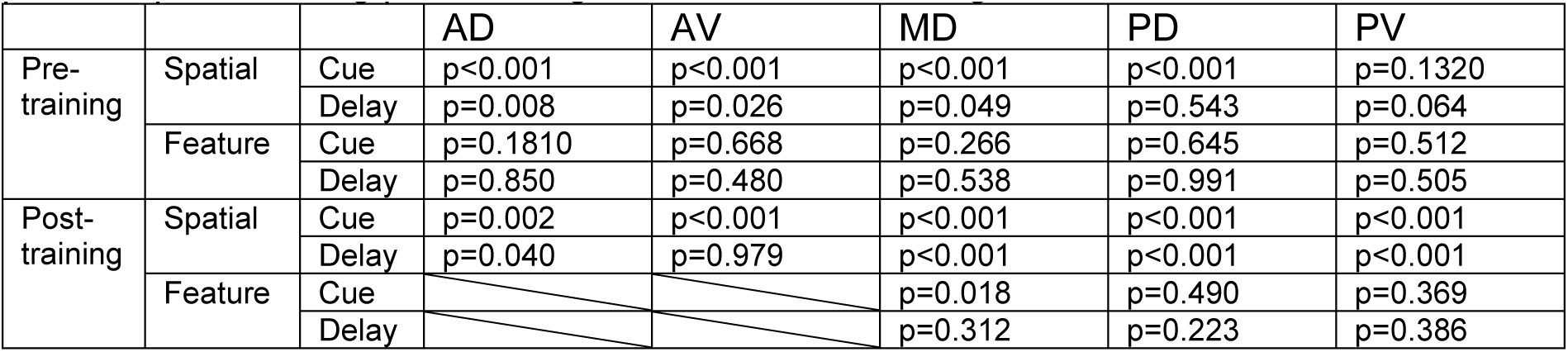
Nonparametric test statistics for order regularity compared to random order from the pre- and post-training phase in Fig.2B, number of shuffling iterations=1000.

**Table S4:** Paired t-test of rotations angles from “within” and “cross” for cue vs sample match, where the sample size n=100.

|  |  | AD | AV | MD | PD | PV |
| --- | --- | --- | --- | --- | --- | --- |
| Spatial | Pre | t(99)=15.7, p<0.001 | t(99)=12.2, p<0.001 | t(99)=8.49, p<0.001 | t(99)=7.64, p<0.001 | t(99)=12.7, p<0.001 |
|  | Post | t(99)=12.9, p<0.001 | t(99)=12.8, p<0.001 | t(99)=7.08, p<0.001 | t(99)=13.1, p<0.001 | t(99)=2.60, p=0.011 |
| Feature | Pre | t(99)=12.9, p<0.001 | t(99)=21.1, p<0.001 | t(99)=4.35, p<0.001 | t(99)=7.58, p<0.001 | t(99)=2.51, p=0.014 |
|  | Post |  |  | t(99)=4.86, p<0.001 | t(99)=3.87, p<0.001 | t(99)=5.07, p<0.001 |

**Table S5:** Paired t-test of rotations angles from “within” and “cross” for sample-match vs sample-nonmatch, where the sample size n=100.

|  |  | AD | AV | MD | PD | PV |
| --- | --- | --- | --- | --- | --- | --- |
| Spatial | Pre | t(99)=28.3,p<0.001 | t(99)=22.5,p<0.001 | t(99)=8.97,p<0.001 | t(99)=17.8,p<0.001 | t(99)=42.0,p<0.001 |
|  | Post | t(99)=31.5,p<0.001 | t(99)=40.8,p<0.001 | t(99)=52.2,p<0.001 | t(99)=10.8,p<0.001 | t(99)=29.0p<0.001 |
| Feature | Pre | t(99)=30.4,p<0.001 | t(99)=22.7,p<0.001 | t(99)=21.7,p<0.001 | t(99)=45.3,p<0.001 | t(99)=40.5,p<0.001 |
|  | Post |  |  | t(99)=14.5,p<0.001 | t(99)=8.1,p<0.001 | t(99)=17.2,p<0.001 |

## Notes

### Competing Interest Statement

The authors have declared no competing interest.

### Summary of Updates

Provided more in-depth results; updated main and supplementary figures; expanded on methods.

